# Objects sharpen visual scene representations: evidence from MEG decoding

**DOI:** 10.1101/2023.04.06.535903

**Authors:** Talia Brandman, Marius V. Peelen

## Abstract

Real-world scenes consist of objects, defined by local information, and scene background, defined by global information. While objects and scenes are processed in separate pathways in visual cortex, their processing interacts. Specifically, previous studies have shown that scene context makes blurry objects look sharper, an effect that can be observed as a sharpening of object representations in visual cortex from around 300 ms after stimulus onset. Here, we use MEG to show that objects can also sharpen scene representations, with the same temporal profile. Photographs of indoor (closed) and outdoor (open) scenes were blurred such that they were difficult to categorize on their own but easily disambiguated by the inclusion of an object. Classifiers were trained to distinguish MEG response patterns to intact indoor and outdoor scenes, presented in an independent run, and tested on degraded scenes in the main experiment. Results revealed better decoding of scenes with objects than scenes alone and objects alone from 300 ms after stimulus onset. This effect was strongest over left posterior sensors. These findings show that the influence of objects on scene representations occurs at similar latencies as the influence of scenes on object representations, in line with a common predictive processing mechanism.

## Introduction

We need only a glance at a new place to determine whether it is a field or a forest, a street or a playground (Potter, 1976; Fei-Fei et al., 2007), while at the same time we rapidly categorize individual objects within that scene (Thorpe et al., 1996; Fei-Fei et al., 2007). This is an example of how we naturally distinguish between the recognition of scenes, informed by global information, and of objects, informed by local information. In line with this intuitive distinction, neuroimaging studies have provided evidence for a neural distinction between the visual processing of scenes and objects (Epstein, 2014). Specifically, scenes and objects activated distinct visual cortex regions in functional magnetic resonance imaging (fMRI) studies, with objects preferentially activating the lateral occipital cortex (LO) and posterior fusiform gyrus (pFs; Grill-Spector and Malach, 2004) and scenes preferentially activating the parahippocampal place area (PPA; Epstein and Kanwisher, 1998), the retrosplenial complex (RSC; Maguire et al., 2001), and the occipital place area (OPA) near the transverse occipital sulcus (TOS; Grill-Spector, 2003). Transcranial magnetic stimulation (TMS) studies have provided causal evidence to support these findings, showing that stimulation of the scene-selective OPA selectively impaired scene recognition while stimulation of the object-selective LO selectively impaired object recognition (Mullin and Steeves, 2011; Dilks et al., 2013; Ganaden et al., 2013; Wischnewski and Peelen, 2021a).

In addition to evidence for the separate processing of scenes and objects, there is also mounting evidence that their processing interacts. A large behavioral literature has focused on the effects of scene context on object recognition, showing that object recognition is more accurate when objects appear in a congruent as compared with an incongruent context (Biederman et al., 1982; Bar, 2004; Davenport and Potter, 2004; Oliva and Torralba, 2007; Munneke et al., 2013; Võ et al., 2019). While it was initially unclear to what extent these effects occurred at the perceptual and/or decisional level (Henderson and Hollingworth, 1999), recent studies have provided evidence that scene context can, in some cases, perceptually sharpen the representation of objects (Rossel et al., 2022, 2023). At the neural level, fMRI studies have shown that scene context modulates the representation of objects in visual cortex. Specifically, the representations of ambiguous (degraded) objects in object-selective LO and pFs became more similar to the representations of the corresponding intact objects when the ambiguous objects were presented in their original scene context (Brandman and Peelen, 2017). A magnetoencephalography (MEG) study further showed that this scene-based sharpening of object representations occurred at around 300 ms after stimulus onset (Brandman and Peelen, 2017). Finally, it was shown that LO activity at this time causally contributed to scene-based object recognition, such that TMS over LO at 260-300 ms after stimulus onset selectively impaired scene-based object recognition, but not isolated-object recognition (Wischnewski and Peelen, 2021b).

One interpretation of these findings is that rapidly processed scene “gist”, based on coarse (low spatial frequency) information, activates a scene schema that then facilitates object processing (Schyns and Oliva, 1994; Bar, 2004; Oliva and Torralba, 2007). This interpretation implies that scene-object interactions are unidirectional, with scene context informing object processing but not necessarily vice versa. Interestingly, however, there is now also evidence for the reverse influence, with objects facilitating scene processing. For example, behavioral studies have found that scene recognition was more accurate for scenes shown together with a semantically congruent (vs incongruent) object (Davenport and Potter, 2004; Davenport, 2007; Leroy et al., 2020) and, like scene-to-object influences, such object-to-scene congruency effects were also found for briefly presented and masked images of scenes (Joubert et al., 2007; Furtak et al., 2022). Finally, objects have also been shown to modulate scene representations in scene-selective regions of visual cortex. Specifically, the representations of ambiguous (degraded) scenes in left OPA and left PPA became more similar to representations of the corresponding intact scenes when the ambiguous scenes were presented together with a congruent object (Brandman and Peelen, 2019). To our knowledge, the time course of this object-based sharpening of scene representations has not been investigated. Revealing this time course would allow for comparing the time courses of scene-to-object and object-to-scene influences.

To fill this gap, here we used MEG to investigate when the representation of scenes is modulated by object presence. The design and analysis approach closely followed that of our previous fMRI study (Brandman and Peelen, 2019). Specifically, we measured the multivariate representations of scene category evoked by degraded scenes with objects, degraded scenes alone, and objects alone. In a separate intact-scenes experiment used for classifier training, participants viewed indoor (closed) and outdoor (open) intact scene images, presented without the foreground object. We used cross-decoding classification of scene category (indoor/outdoor) to compare the multivariate response patterns evoked by intact scenes to those evoked by the main experiment stimuli. This allowed us to measure the contribution of the object to the neural representation of scene category in degraded scenes, in order to characterize the temporal dynamics of object-based facilitation of visual scene processing.

Based on our previous study, showing that object representations are disambiguated by scene context from around 300 ms after stimulus onset (Brandman and Peelen, 2017), we hypothesized that scene representations would similarly be disambiguated by objects at 300 ms after stimulus onset.

## Materials & Methods

### Participants

Twenty-eight healthy participants (13 male, mean 26 years ± 4 SD) took part in the study. All participants had normal or corrected to normal vision and gave informed consent. Sample size was chosen to match that of our previous studies using similar MEG decoding methods (Brandman and Peelen, 2017; Brandman et al., 2020). No subjects were excluded. All procedures were approved by the ethics committee of the University of Trento.

### Stimuli

The stimulus set was the same as the set used in the corresponding fMRI study (Brandman and Peelen, 2019), consisting of degraded scenes that were perceived as ambiguous on their own but that were easily categorized when presented with an object (**Figure 1A**). Scenes were degraded in Adobe Photoshop by applying radial blur and reducing image contrast. Both indoor and outdoor scenes included a mixture of animate and inanimate objects of various categories, and did not contain other objects contextually associated with the foreground objects. The main experiment stimuli consisted of 30 indoor and 30 outdoor scene photographs with one dominant foreground object, presented in 3 conditions (total 180 images): scene with object, scene alone, or object alone on a uniform gray background of mean luminance of the original background (see examples in **Figure 1A**; for the full stimulus set, see https://www.ncbi.nlm.nih.gov/pmc/articles/PMC6033316/bin/NIHMS76424-supplement-Supplementary_Figure.pdf, and for details on stimulus generation, see Brandman and Peelen, 2019). To avoid familiarity effects passing from scenes with objects to scenes alone, the stimulus set was split in two, such that different scenes were presented for degraded scenes with objects and for degraded scenes alone within a given subject, counterbalanced across subjects (Brandman and Peelen, 2019). The pattern localizer included the 60 scenes from the main experiment (with different cropping), and an additional set of 60 new scenes that were matched for category and sub-category of the main experiment set, in high resolution. Importantly, the intact scenes never included the foreground object. All stimuli were presented centrally, at a visual angle of 6x6 degrees (400x400 pixels).

**Figure 1:**
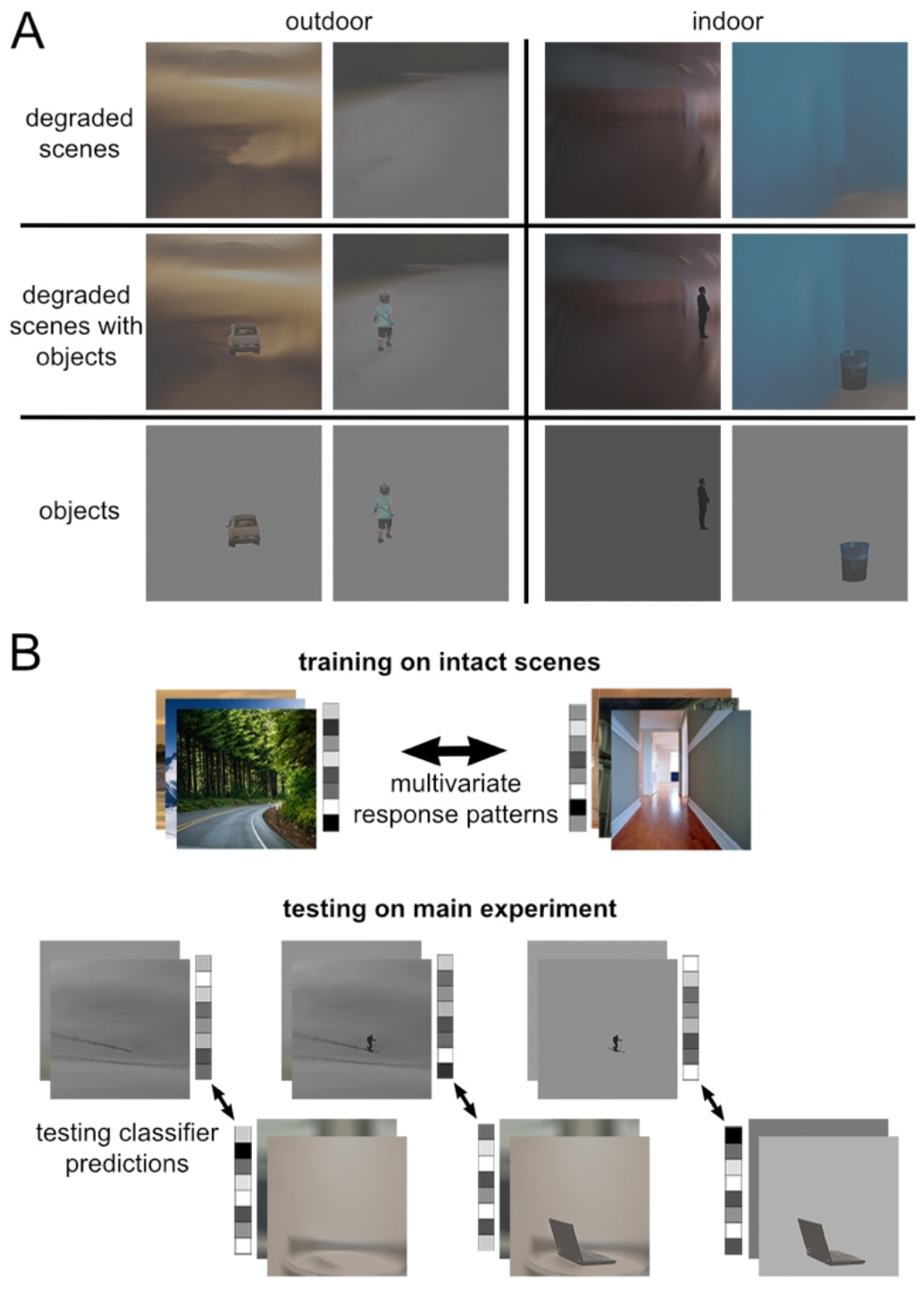
Experimental approach. (A) Sample stimuli used in the main experiment across 6 conditions: indoor/outdoor X degraded scenes, degraded scenes with objects, objects on uniform gray background of mean luminance of the original background. For the full stimulus set, see Brandman and Peelen (2019). (B) Cross-decoding of scene category: we trained a linear classifier on the multivariate response pattern across MEG sensors to indoor and outdoor intact scene photographs, and tested classifier predictions of scene category based on the response patterns for each of the main experiment conditions.

### Procedure

On each trial, participants viewed a 500 ms fixation screen followed by the stimulus briefly presented for 83 ms, and a jittered inter-trial-interval (ITI) between 1200-1600 ms (mean ITI 1400 ms). Throughout all runs participants performed a 1-back task in which they pressed a button each time an image appeared twice in a row. Performance on this task was highly accurate (mean accuracy = 97%, SD = 5%). Every 33 trials they were presented with their response accuracy for 5 s, and once in the middle of the run they received an additional 8 s fixation break. The main experiment consisted of 4 runs of ∼426 s duration, including 18 one-back target trials and 30 trials per condition: indoor/outdoor x scene/object/scene-with-object (198 trials/run). The intact scene experiment consisted of 4 runs of ∼285 s duration, including 12 1-back trials and 30 trials per condition: indoor/outdoor x old/new (132 trials/run). This resulted in 120 trials per condition in the main experiment, and 120 trials per condition in the intact scene experiment. Trials were randomly intermixed within each run.

### Data Acquisition and Preprocessing

Electromagnetic brain activity was recorded using an Elekta Neuromag 306 MEG system, composed of 204 planar gradiometers and 102 magnetometers. Signals were sampled continuously at 1000 Hz and band-pass filtered online between 0.1 and 330Hz. Offline preprocessing was done using the Elekta MaxFilter/MaxMove software, MATLAB (RRID:SCR_001622) and the FieldTrip analysis package (RRID:SCR_004849). Spatiotemporal signal space separation (tSSS) was performed to reduce noise originating from external (non-brain) signals, as well as noise produced by head motion, by exploiting certain properties of the solution to the Maxwell equations (Taulu and Simola, 2006), as implemented by the Elekta MaxFilter/MaxMove software. Data were then demeaned, detrended, down-sampled to 100 Hz and time-locked to visual onset. The data were averaged across trials of the same exemplar across runs (excluding 1-back trials, which were discarded), resulting in a total of 90 unique test trials throughout the main experiment (15 per condition), and 120 unique train/test trials throughout the intact scenes experiment (30 per condition). Except for the 1-back trials, no trials were excluded and no additional artefact removal methods were used.

### Multivariate Analysis

Multivariate analysis was performed using the CoSMoMVPA toolbox (Oosterhof et al., 2016) (RRID:SCR_014519). Analysis followed a similar procedure as in our fMRI study using the same approach (Brandman and Peelen, 2019), in which a cross-decoding algorithm was trained on scene-category classification of intact scenes (without foreground objects) and tested on degraded scenes, degraded scenes with objects, and objects alone. Decoding was performed across the 24 left-hemisphere posterior magnetometers of each participant between 0 and 500 ms. We focused on left posterior channels based on our previous fMRI findings of a left-lateralized effect in scene-selective visual areas using the same decoding approach (Brandman and Peelen, 2019). Magnetometers were used as these gave the most reliable classification in previous work using similar decoding methods (Kaiser et al., 2016; Brandman et al., 2020). Prior to decoding, temporal smoothing was applied by averaging across neighboring time-points at a distance of 2 (20 ms) on each side. An LDA classifier discriminated between response patterns to indoor vs. outdoor scenes. The covariance matrix was regularized by adding the identity matrix scaled by 0.01 of the mean of the diagonal (as implemented in CoSMoMVPA). The decoding approach is illustrated in **Figure 1B**. First, decoding of intact scene category was measured within the pattern localizer, by training on old-scene trials (i.e., scenes included in the main-experiment set, without the foreground object; 60 samples), and testing on new-scene trials (60 samples). Next, cross-decoding was achieved by training on all conditions of the pattern localizer (120 samples), and testing on each of the main-experiment conditions (scene, scene-with-object, object; 30 samples each). Decoding was performed for every possible combination of training and testing time-points between 0 and 500 ms, resulting in a 50 x 50 matrix of 10 ms time-points, for each of the tested conditions, per subject. In addition, to generate a measure of same-time cross-decoding, decoding accuracy of each time-point along the diagonal of the matrix was averaged with its neighboring time-points at a radius of 2 (20 ms in every direction).

### Significance testing

Classification significance against chance (accuracy of 0.5) was tested on decoding accuracies across the time-by-time matrix. Significance was tested for each time-point by computing random-effect temporal-cluster statistics corrected for multiple comparisons, as implemented in CoSMoMVPA. This was accomplished via t-test computation over 1000 permutation iterations, in which the sign of samples was randomly flipped (over all features) after subtracting the mean, and using threshold free cluster enhancement (TFCE) as cluster statistic, with a threshold step of 0.1. Significance was determined by testing the actual TFCE image against the maximum TFCE scores yielded by the permutation distribution (TFCE, *p* < 0.05) (Smith and Nichols, 2009). Significant above-chance decoding of scene category in the intact-scenes experiment was tested across the entire time-by-time matrix. Significant time-points were then used to define a temporal mask for cross-decoding significance testing (**Figure 2**). Paired differences between main experiment conditions were tested along the same-time (i.e., of training and testing) cross-decoding accuracies, similarly using TFCE and the temporal mask.

**Figure 2:**
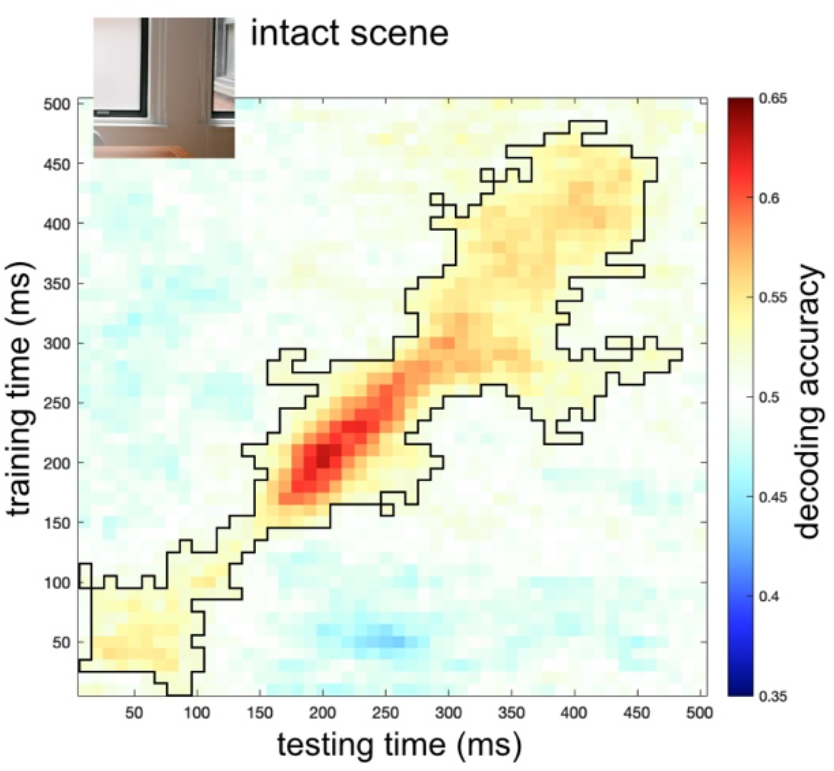
Decoding of scene category from activity evoked by intact images. Decoding accuracies from a classifier trained to distinguish indoor vs outdoor intact scenes (**Figure 1B**) across two stimulus sets. Matrices represent the decoding accuracy across a time x time space from stimulus onset to 500 ms, averaged across participants. Black outlines denote clusters for which decoding was significantly above chance (classifier accuracy 0.5) at TFCE, p < 0.05.

### Searchlight Analysis

The same cross-decoding method described above was applied in a searchlight approach (Kriegeskorte et al., 2006; Kaiser et al., 2016), across the entire scalp of each participant, as implemented in CoSMoMVPA. Searchlight was performed using all magnetometers (similar clusters were observed when using both magnetometers and gradiometers). For each timepoint, the searchlight analysis was performed separately for scenes with objects and scenes alone, across neighborhoods of 15 channels (Kaiser et al., 2016), resulting in an accuracy score for each channel, condition, and timepoint. Thereafter, decoding accuracies were averaged into time-clusters of 50 ms each, resulting in 10 decoding maps between 0 and 500 ms for each condition. Within each time-cluster, significant object-based scene facilitation was tested by the difference between scenes with objects and scene alone, using the same permutation and TFCE procedures as described above, applied to the spatial distribution of decoding accuracies across channels.

## Results

We tested above-chance classification of superordinate scene category using a cross-decoding approach (**Figure 1B**). In the main analyses, classifiers were trained to discriminate indoor versus outdoor intact scenes (without foreground objects), and then tested on scene category discrimination of degraded scenes with objects, degraded scenes alone, and objects alone on a uniform gray background. In our previous fMRI study using the same stimuli and analysis approach, we found a strongly lateralized effect in scene-selective areas, with object-based scene facilitation restricted to the left hemisphere (Brandman and Peelen, 2019). Therefore, our analyses focused on left hemisphere sensors. In addition, to examine the spatial distribution of scene decoding across the entire scalp, we ran a searchlight analysis through all MEG sensors.

### Decoding scene category from intact scenes

In a first analysis, we decoded scene category (indoor/outdoor) from intact scenes. The significant cluster in the time x time decoding matrix spanned most of the diagonal, peaking at around 200 ms after stimulus onset (**Figure 2**). These results show that MEG response patterns to indoor and outdoor scenes reliably differed at multiple stages of visual processing.

### Decoding scene category from degraded scenes

In the main analyses, classifiers were trained on intact scenes (without foreground objects) and tested on degraded scenes alone, degraded scenes with objects, and objects alone. Cross-decoding accuracies are presented in **Figure 3**. Scene category could be reliably decoded from degraded scenes alone between ∼200-300 ms (**Figure 3A**), and from degraded scenes with objects between ∼250-400ms (**Figure 3B**). By contrast, scene category could not be decoded from objects alone at any time point (**Figure 3C**). These results show that superordinate scene category could be extracted from degraded scenes, particularly when an object was present. Importantly, the object alone could not explain this decoding.

**Figure 3:**
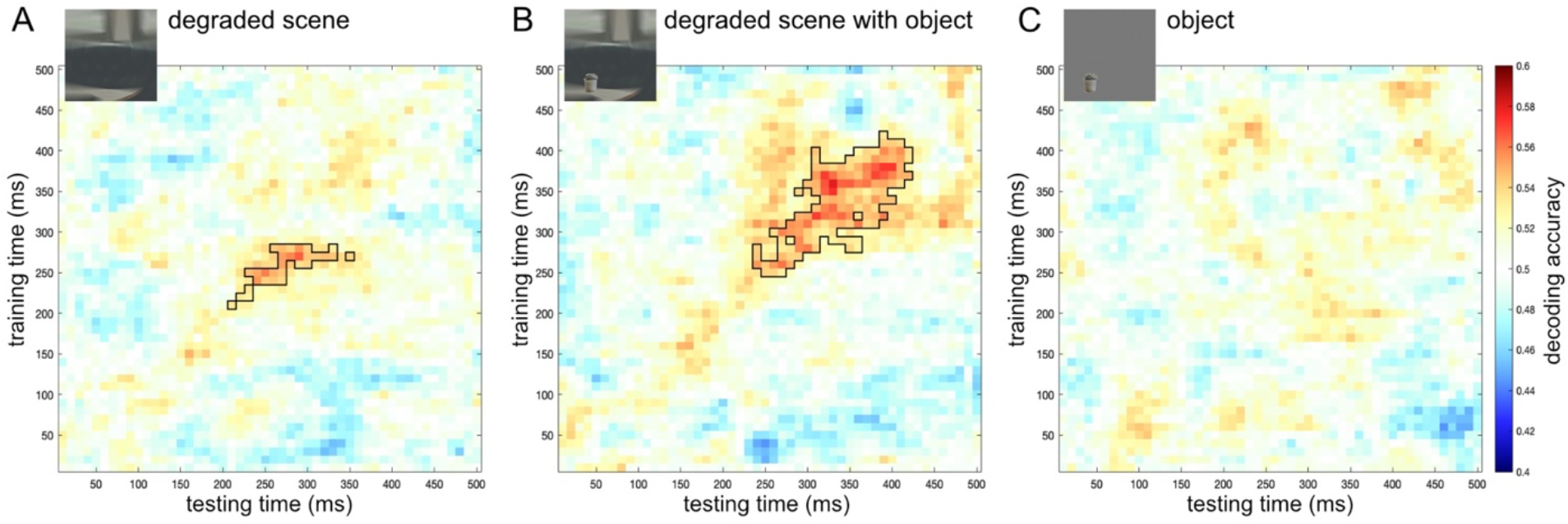
Cross-decoding of scene category. Decoding accuracies from a classifier trained to distinguish indoor vs outdoor intact scenes (**Figure 1B**) and tested on (A) degraded scenes, (B) degraded scenes with objects, and (C) objects alone. Matrices represent the decoding accuracy across a time x time space from stimulus onset to 500 ms, averaged across participants. Black outlines denote clusters for which decoding was significantly above chance (classifier accuracy 0.5) at TFCE, p < 0.05.

### Object-based scene facilitation

To measure object-based scene facilitation, we compared the cross-decoding accuracies of degraded scenes with and without objects (and objects alone), along similar training and testing times from stimulus onset (**Figure 4**). Significant object-based scene facilitation was found around 320 ms after stimulus onset. Specifically, decoding accuracies were significantly higher for degraded scenes with objects than for degraded scenes alone between 310-330 ms, and were also higher for degraded scenes with objects than for objects alone between 320-350 ms. These results provide evidence that objects facilitated scene processing.

**Figure 4:**
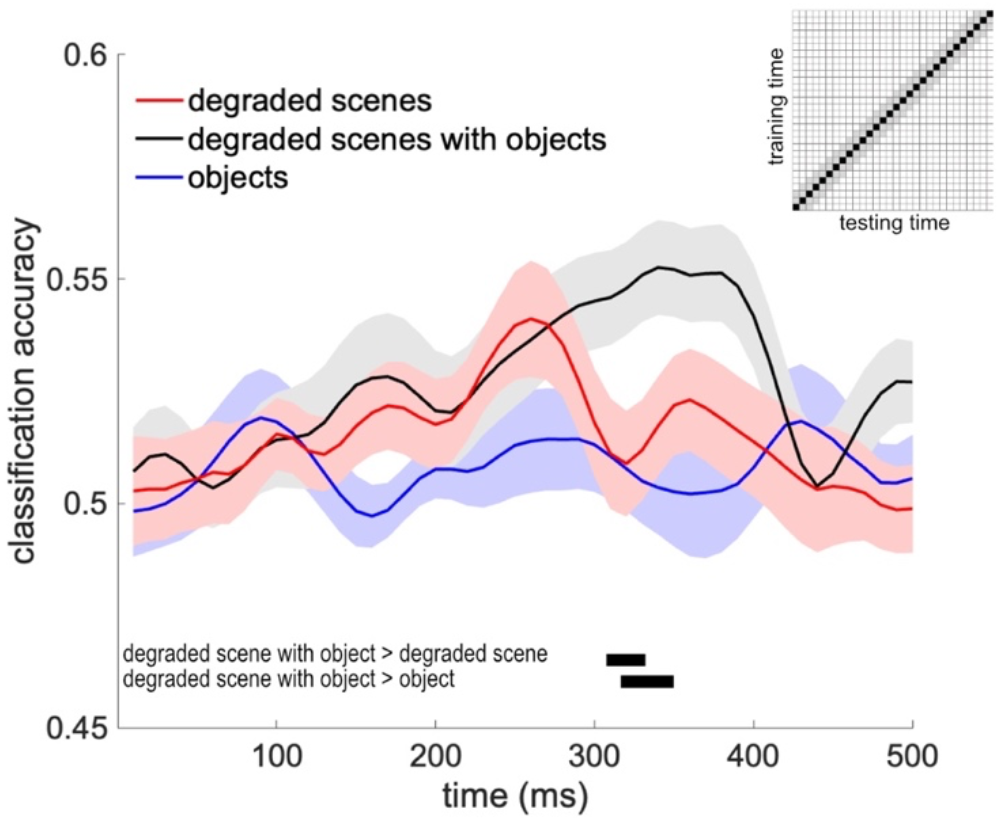
Object-based scene facilitation. Cross-decoding accuracies along the time diagonal (matched training and testing times) of the matrices presented in **Figure 3**. Data are represented as mean ± SEM across subjects. Black-filled rectangles denote significant differences between conditions at TFCE, p < 0.05.

### The spatial distribution of object-based scene facilitation

In the final analysis, we measured cross-decoding accuracies for degraded scenes with and without objects across the entire scalp in a searchlight procedure (similar results were obtained for the less controlled comparison of degraded scenes with objects versus objects alone). Searchlight results are presented in **Figure 5**. Significant object-based scene facilitation was found between 300-350 ms, mostly in left posterior sensors (**Figure 5A**). In these sensors, the average neighborhood decoding accuracy was higher for degraded scenes with objects than for degraded scenes alone. Scene category could be decoded from both degraded scenes (**Figure 5B**) and degraded scenes with objects (**Figure 5C**) from around 200 ms. These results show that object-based facilitation followed the initial representation of scene category extracted from degraded scenes.

**Figure 5:**
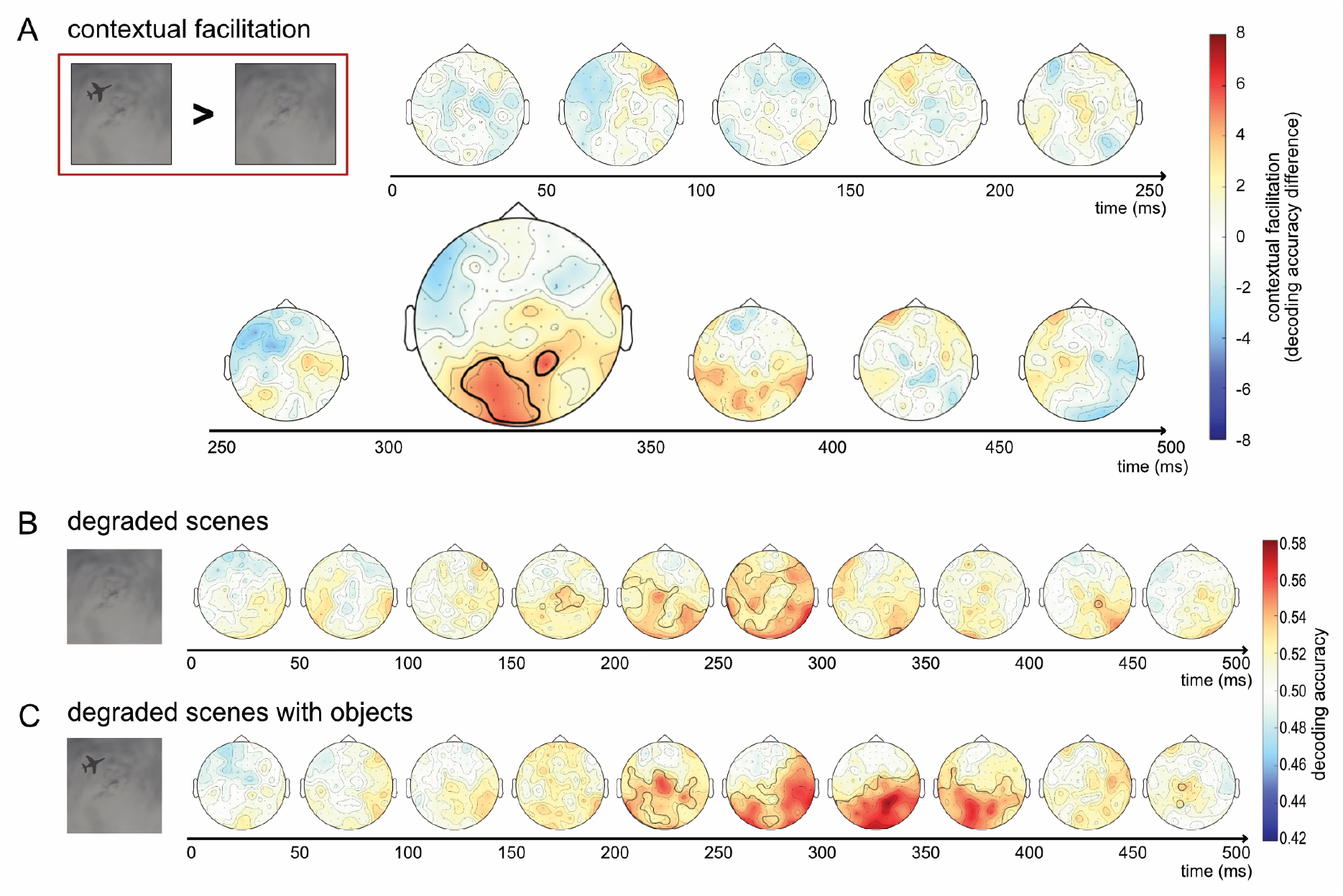
Searchlight of object-based scene facilitation. (A) Difference in cross-decoding accuracies for degraded scenes with objects versus degraded scenes alone. Black outlines denote significant object-based scene facilitation, measured by the difference between degraded scenes with and without objects, TFCE p < 0.05. (B) Decoding accuracy for degraded scenes alone and (C) degraded scenes with objects. Black outlines denote significant above-chance decoding, TFCE p < 0.05. Decoding results were averaged across neighborhoods of 15 sensors in a searchlight procedure, averaged across 50 ms time-clusters, and averaged across participants.

## Discussion

The current results provide evidence that objects sharpen the representation of ambiguous scenes from 300 ms after stimulus onset. At this time point, the multivariate response patterns evoked by the degraded (ambiguous) scene, when presented with an object, became more similar to the multivariate response patterns evoked by the corresponding scene when presented intact and fully visible. Importantly, the intact scenes used for classifier training did not contain the foreground objects that disambiguated the degraded scenes. As such, the visual processing of the foreground objects themselves could not account for the increased decoding; if anything, the intact scenes used for training the classifier were visually more similar to the degraded scenes without objects than the degraded scenes with objects. Indeed, in our previous fMRI study, using the same stimuli and analysis approach, decoding in early visual cortex was higher for the degraded scenes without objects than for the degraded scenes with objects. Therefore, we interpret the increased decoding for degraded scenes with objects as reflecting a sharpening of the representation of the background scene. In our previous fMRI study, we observed a sharpening of scene representations in the left OPA and left PPA (Brandman and Peelen, 2019). This accords well with the current whole-brain searchlight results, which showed that the strongest object-based facilitation in decoding was observed over left posterior sensors (**Figure 5A**). Combining the results of both studies supports the conclusion that objects sharpen representations in left scene-selective visual cortex from 300 ms after stimulus onset.

What scene properties are disambiguated by objects? Classifiers were trained on distinguishing indoor versus outdoor scenes, categories that differ at multiple levels, from low-level features to their semantic category and the actions they afford (Greene and Hansen, 2020). Indeed, classifiers that were trained and tested on (different exemplars of) intact scenes showed above-chance decoding across a broad time window (50-450 ms after stimulus onset). Decoding along this time window likely reflects different stages of scene processing, as also suggested by the limited generalization across time (**Figure 2**; King and Dehaene, 2014). The presence of objects could potentially lead to a sharpening of all these levels of representation through feedback within the scene pathway (Peelen et al., 2023). Notably, the strongest decoding of ambiguous scenes with objects was observed along the diagonal of the time x time matrix from 250 ms after scene onset (**Figure 3**), suggesting that relatively late-stage representations evoked by the intact scenes were disambiguated by the objects. One possibility is that the object (e.g., an airplane) helped to recognize the ambiguous scene at the basic level (e.g., sky) which then led to activation of superordinate category (indoor vs outdoor) properties. These properties include spatial layout (open vs closed), which is represented around 250 ms after stimulus onset (Cichy et al., 2017) and is also represented in OPA and PPA (Kravitz et al., 2011; Park et al., 2011). Accordingly, our findings may reflect the disambiguation of the spatial layout of the scene. However, future studies are needed to systematically manipulate clear and ambiguous scene properties to test this more conclusively. For example, it would be interesting to test whether and when disambiguation is observed at the basic level (e.g., distinguishing between two outdoor scene categories). Relatedly, studies could investigate whether the timing of the facilitation depends on the task relevance of the categorization, e.g., whether objects facilitate scene representations more quickly when the decoded dimension is relevant for the task.

The current study complements our previous MEG study investigating the reverse effect: the sharpening of object representations by scene context (Brandman and Peelen, 2017). In that study, we found that the decoding of object category (animate vs inanimate) increased for ambiguous objects presented within their original scene context, both when compared with the same objects outside of scene context and when compared with the scene context alone. Almost identical to the current results, this increase was observed from 320-340 ms after stimulus onset. Together, these studies show that the interaction between scene and object processing is bidirectional, without a clear temporal asymmetry. This is in line with behavioral studies showing bidirectional interactions between objects and scenes in recognition and categorization tasks (Davenport and Potter, 2004; Davenport, 2007; Joubert et al., 2007; Leroy et al., 2020; Furtak et al., 2022). Furthermore, studies using free-report paradigms have shown that participants are equally likely to report scene and object features for scenes presented very briefly. Rather than demonstrating a categorical advantage (e.g., scenes before objects), participants report lower-level features of both scenes and objects for short presentation times and increasingly higher-level features (e.g., semantic categories) of both scenes and objects for longer presentation times (Fei-Fei et al., 2007; Chuyin et al., 2022). Nevertheless, while interactions between objects and scenes are bidirectional and do not appear to show a clear temporal asymmetry, in daily life it is likely that scenes exert a stronger influence on object processing than vice versa, considering that scene information is usually less ambiguous and more stable over time than object information.

Our findings can be explained within a general predictive processing framework, in which recurrent feedforward/feedback loops in the cortex serve to integrate top-down contextual priors and bottom-up observations so as to implement concurrent probabilistic inference along the visual hierarchy (Lee and Mumford, 2003). Our results support this interpretation by showing that scene category is initially represented for degraded scenes with and without objects, with objects subsequently boosting scene category information (**Figure 5**). Contextual matching of bottom-up input and top-down expectations has also been proposed to account for object-based scene processing (Bar and Ullman, 1996; Bar and Aminoff, 2003; Bar, 2004; Mudrik et al., 2014). Interestingly, scenes and objects are initially processed in separate pathways that are not hierarchically related to each other. This raises the question of how information from one pathway reaches the other pathway. One possibility is that object and scene pathways interact at higher (e.g., categorical) stages, with information then feeding back within each pathway, leading to perceptual sharpening (Peelen et al., 2023). For example, recognizing a scene as a road could lead to the disambiguation of a car-sized blob as a car, which would then lead to the disambiguation of specific car features (e.g., the taillight).

If interaction takes the form of a representational sharpening loop, converging with the accumulation of sufficient evidence, we may ask how unambiguous stimuli are processed. Many objects and scenes we see in everyday contexts already provide sufficient feed-forward information to identify them without the need for contextual integration. Current data suggests that contextual sharpening occurs on a need-to-know basis, i.e., when feed-forward intrinsic information is insufficient, informing multiple plausible representations, extrinsic information is gathered via contextual processing until a unique representation emerges. In line with this notion, behavioral data revealed that the facilitating effect of the scene on object detection was reduced for intact objects, and that object ambiguity was correlated with the effectiveness of contextual facilitation (Brandman and Peelen, 2017). Other studies have shown that scene-based expectations benefit the processing of scene-congruent objects when the objects are ambiguous, but benefit the processing of scene-incongruent objects when there is no ambiguity (Spaak et al., 2022; Rossel et al., 2023). Finally, an electrophysiology (EEG) study showed that ERP differences in response to congruent and neutral contexts were larger for ambiguous than for unambiguous objects (Dyck and Brodeur, 2015). While these studies investigated the influence of scene context on object processing, we expect that similar principles apply to the reverse influence, from objects to scenes. It would be interesting to adopt recently developed behavioral paradigms showing perceptual sharpening of object representations (Rossel et al., 2022) to investigate object-based sharpening of scene representations, also as a function of ambiguity (Rossel et al., 2023).

Finally, an important open question is whether the disambiguation we observe is necessary for scene understanding or whether it is epiphenomenal. We have previously used TMS to show that the scene-based sharpening of object representations at 300 ms after stimulus onset is causally involved in object recognition (Wischnewski and Peelen, 2021b). With the information about the timing of the reverse effect, obtained in the current study, a similar TMS study can now be performed, testing whether object-based sharpening of scene representations at 300 ms after stimulus onset is causally involved in scene recognition.

## Conclusion

Our findings support interactive accounts of object and scene processing, whereby the two pathways generate complementary local and global representations of scenes, and dynamically share information across pathways in order to construct the full percept of a scene. The present study shows that the influence of objects on scene representations occurs at similar latencies as the influence of scenes on object representations (Brandman and Peelen, 2017), in line with a common predictive processing mechanism.

## Acknowledgements

The project was funded by the European Union’s Horizon 2020 research and innovation program under the Marie Sklodowska-Curie grant agreement no. 659778 and by the European Research Council (ERC) under the European Union’s Horizon 2020 research and innovation program (grant agreement no. 725970). This manuscript reflects only the authors’ view, and the Agency is not responsible for any use that may be made of the information it contains.

